# Comparison of antimicrobial activity between intraoperatively soaked bacitracin sutures and triclosan coated suture

**DOI:** 10.1101/2020.08.24.264820

**Authors:** Shane T. Musick, Jeremy M. Adkins, Roy Al-Ahmar, Hongwei D. Yu, Anthony M. Alberico

**Author notes:** Corresponding Author (SM). These authors contributed equally to this work (Shane T. Musick, Jeremy M. Adkins). These authors also contributed equally to this work (Roy Al-Ahmar, Hongwei D. Yu, Anthony M. Alberico).

## Abstract

With the easily available option for surgeons to soak their suture in anti-biotic irrigating solution intraoperatively in mind, this study was designed to evaluate the ability of suture soaked in bacitracin irrigating solution to inhibit the growth of *Staphylococcus aureus* and *Methicillin-resistant Staphylococcus aureus*. Using standard experimental procedure, sterile suture was soaked in Bacitracin suture, and dried for 10 minutes or 6 hours, incubated for 24 h on inoculated plates, and examined for zone of inhibition around the suture. This was compared to control unsoaked suture and antimicrobial suture (AMS) currently on the market to determine if the minor intra operative procedural change of placing suture in antibiotic irrigation solution instead of on the sterile table could confer some antimicrobial activity. The study found the Bacitracin soaked suture (BSS) consistently inhibited the growth of the test organisms. For both organisms, the BSS exhibited a significantly larger zone of inhibition compared to the unsoaked control suture. However, the AMS currently on the market exhibited a larger zone of inhibition compared to the BSS. Placing sutures in a bacitracin irrigation solution intraoperatively instead of directly on the sterile table can achieve some of the in vitro antimicrobial effect seen from AMS currently on the market. This may result in reduced rates of SSIs and associated costs without major procedural change and at reduced overhead.

## Introduction

Surgical site infections (SSIs) can be a complication of any surgical procedure and, as such, antimicrobial suture (AMS) represents a strategy developed in an effort to mitigate this risk. Some studies have suggested that AMS reduces SSIs in a wide range of procedures (1–9). Yet, others have suggested they have no such role (10, 11). Currently, the role of AMS in reducing SSIs remains controversial. Furthermore, early data has largely focused on their application for gastrointestinal (GI) procedures, but with recent studies supporting their utility, others are exploring their potential utility in ophthalmology and orthopedics (12, 13). Thus, the increasing evidence that AMS play a role in preventing SSIs could eventually lead to higher utilization rates in procedures across specialties.

The aim of this study is to determine whether or not the practice of simply having the OR technician place the closing suture in the bacitracin irrigation solution at the onset of the case rather than setting it on the OR table could result in the conveyance of increased antimicrobial activity to the suture material.

## Methods

### Suture Preparation

A vial of Bacitracin powder (Pfizer Inc., New York) was reconstituted using sterile technique with 0.9% NaCl Irrigation, USP (Aqualite, Illinois) to create a Bacitracin solution with a concentration of 1000 Units/mL to replicate the intraoperative bacitracin irrigation solution concentration typically used intraoperatively. Ethicon (Somerville, NJ) 0 Vicryl was then placed in this Bacitracin solution and soaked for 1 hour to replicate intraoperative soaking time. This is shown in figure 1. This Bacitracin soaked suture (BSS) was then placed in a sterile petri dish at room temperature to dry—one group for 10 minutes to replicate intraoperative drying time outside of the solution and a second group for 6 hours to ensure that any measured antimicrobial activity was provided by the prepared suture, not excess Bacitracin solution.

**Fig 1.**
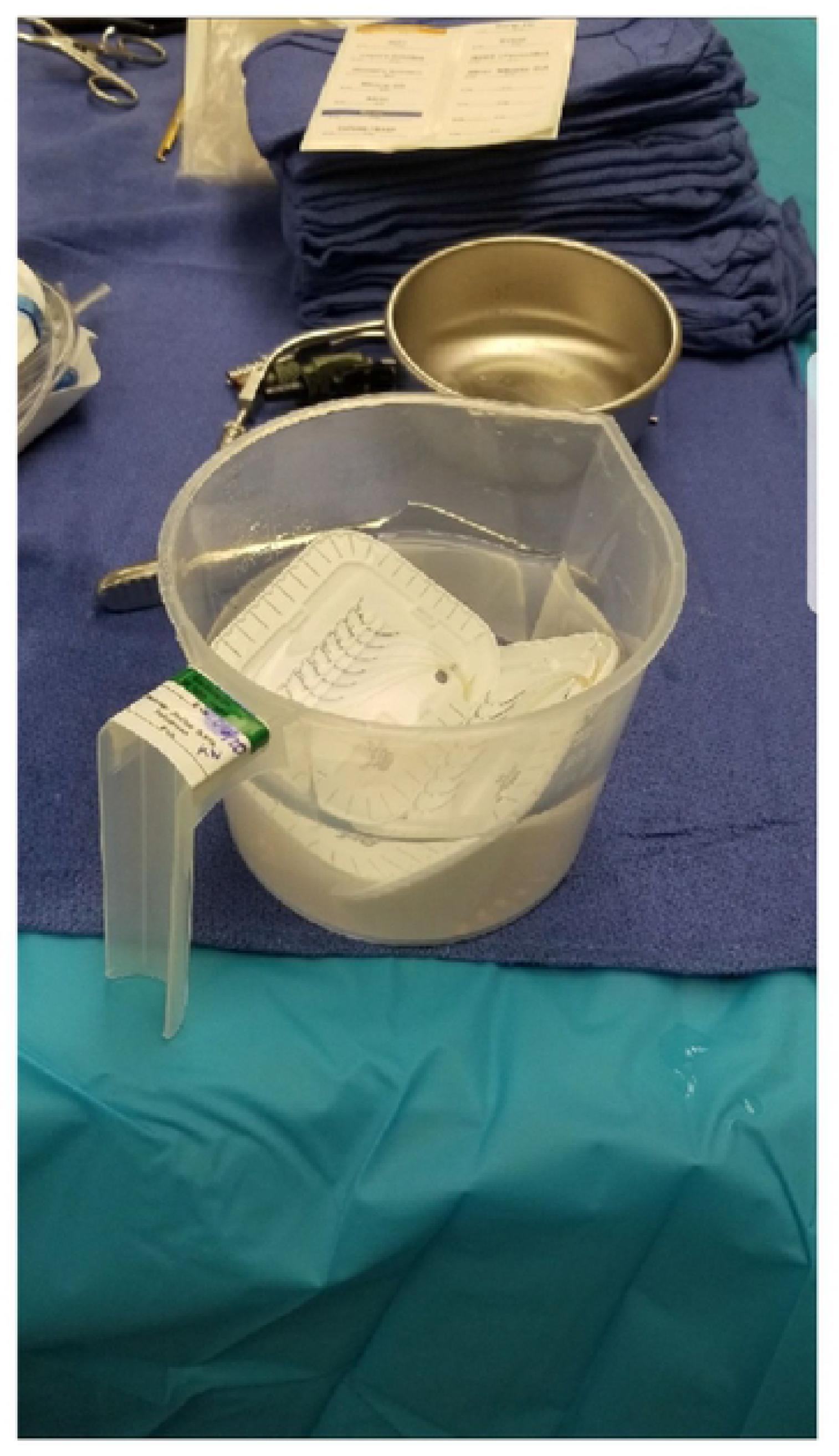
Ethicon 0 VICRYL* sutures soaking in bacitracin irrigation solution intraoperatively. This displays the ease and feasibility of placing suture packs into an antimicrobial solution during the course of the procedure.

### Measuring Antibacterial Activity

Using standard experimental procedure, Staphylococcus aureus (ATCC 6538) and Methicillin Resistant Staphylococcus aureus (MRSA) (N315) were grown in Luria Broth (LB) (Difco) broth at 37°C to a MacFarland Turbidity of 1 (5 ×10^8^ CFU/ml). Subsequently, 100ul of each broth was then inoculated on 27 separate 1.5% agar plates. For each group of 27 agar plates: 3 were plated with one cm of the unsoaked Ethicon 0 VICRYL^®^ Suture as a control, 6 with a 10μl aliquot of the 1000 U/mL Bacitracin solution as an additional control, 6 with one cm sections of BSS dried for 10 minutes, 6 with one cm sections of BSS dried for 6 hours, and 6 with one cm sections of the Ethicon Coated 0 VICRYL^®^ Plus Antibacterial (polyglactin 910). The growth plates were subsequently incubated at 37°C for 24hours, after which they were examined for the presence or absence of a zone of inhibition.

### Statistical Analysis

Statistical analysis was performed using GraphPad Prism 7.02 (GraphPad Software, California). A one-way ANOVA was used followed by Tukey’s multiple comparison test to evaluate the mean difference of the inhibition zone. Student’s t-test was used for individual comparisons. Statistical significance was determined at P < 0.05.

## Results

The size results of the zone of inhibition assays are listed in Table 1. A zone of inhibition was present for all replicates of the Bacitracin solution aliquot, both drying times of the BSS, and Ethicon Coated 0 VICRYL^®^ Plus Antibacterial suture. As expected, the non-antibiotic Ethicon 0 Vicryl failed to exhibit any zone of inhibition. Typical zones of inhibition are shown in Figure 2.

**Table 1.** Comparison of zone of inhibition of *Staphylococcus aureus* and *Methicillin Resistant Staphylococcus aureus (MRSA)*.

|  | 0 VICRYL®<br>(Control) | Bacitracin<br>Solution Aliquot | 10 Minute<br>Dried BSS<sup>a</sup> | 6 Hour<br>Dried BSS<sup>a</sup> | Coated 0 VICRYL®<br>Plus Antibacterial |
| --- | --- | --- | --- | --- | --- |
| Zone of<br>Inhibition v.<br>*S. aureus* | 0 mm | 18.2 mm ± 0.84 | 7 mm ± 1.41 | 7 mm ± 1.79 | 19.5 mm ± 2.07 |
| Zone of<br>Inhibition v.<br>*MRSA* | 0 mm | 15.5 mm ± 1.05 | 5 mm ± 1.22 | 6.83 mm ± 0.75 | 17.67 mm ± 1.37 |

This displays the differing sizes of zones of microbial inhibition across the suture trials. From left to right comparisons - sutures devoid of any bacitracin solution, an aliquot of bacitracin solution itself, 10-minute dried BSS sutures, 6-hour dried BSS sutures, and the pre-coated antibacterial suture.
<sup>a</sup>BSS = Bacitracin solution-soaked Ethicon 0 VICRYL® Suture

**Figure 2:**
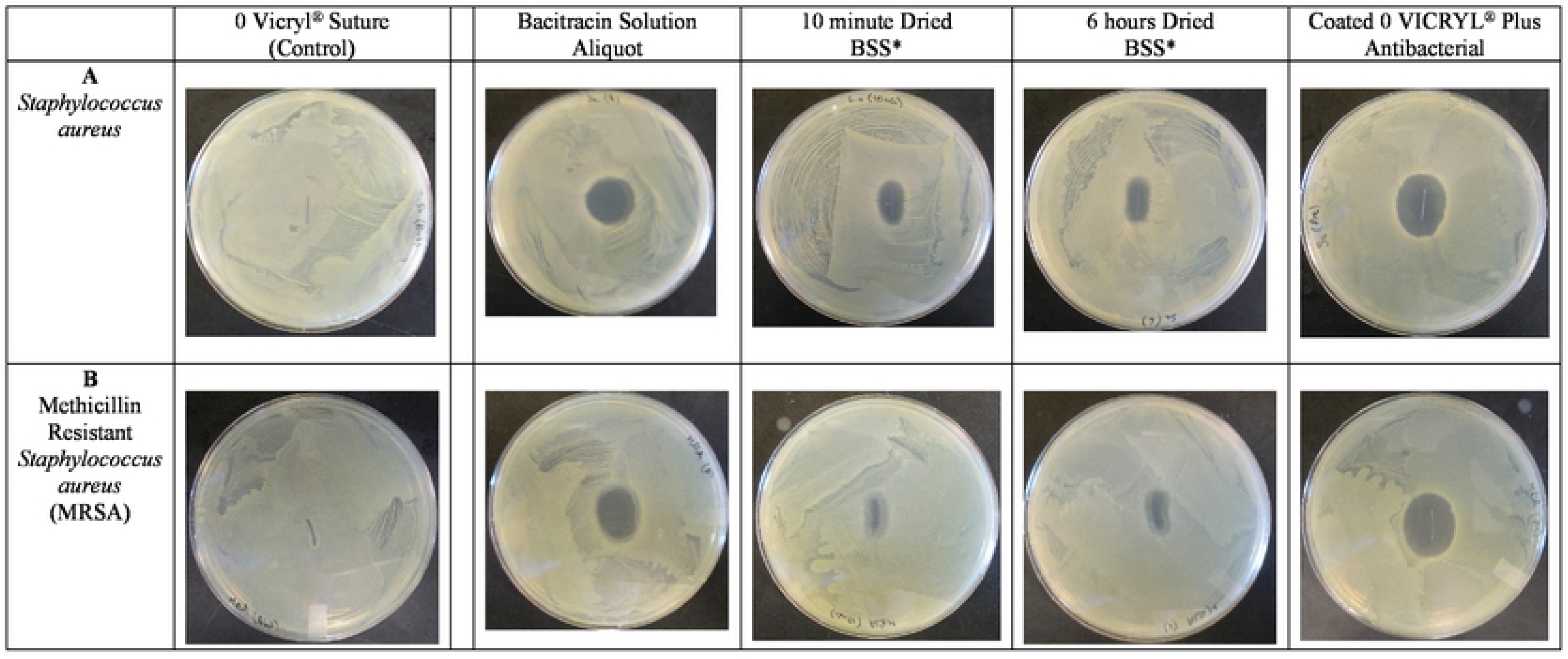
Zone of inhibition petri dish images. **A**: Image showing (from left to right) Staphylococcus aureus in LB broth grown on 1.5% agar plates grown in the presence of 10μl aliquot of 1000U/ml Bacitracin solution, 0 Vicryl® Suture, BSS* dried for 10 minutes, BSS* dried for 6 hours, and Coated 0 VICRYL® Plus Antibacterial after incubation at 37oC for 24hours. **B**: Image showing (from left to right) MRSA in LB broth grown on 1.5% agar plates grown in the presence of 10μl aliquot of 1000U/ml Bacitracin solution, 0 Vicryl® Suture, BSS* dried for 10 minutes, BSS* dried for 6 hours, and Coated 0 VICRYL® Plus Antibacterial after incubation at 37°C for 24hours.

Using a one-way ANOVA, and student’s t-test, the average zone of inhibition was compared between all groups of the study. There were no significant differences between the different drying times using BSS. There were no significant differences detected between Ethicon Coated 0 VICRYL® Plus Antibacterial suture and the Bacitracin solution aliquot (p=0.58). For S. aureus and MRSA, both drying times of the BSS exhibited a significantly larger zone of inhibition than the control unsoaked Ethicon 0 Vicryl (p < 0.0001). However, for S. *aureus* and MRSA, the Ethicon Coated 0 VICRYL^®^ Plus Antibacterial suture and Bacitracin solution aliquot had significantly larger zone of inhibitions than both drying times of the BSS (p < 0.0001). Overall, these results demonstrate that both drying times are similarly effective, suggesting feasibility of use throughout the entirety of a procedure. The BSS displays robust and statistically significant zones of inhibition, indicating antibacterial activity. The zones of inhibition of the Ethicon Coated 0 VICRYL^®^ Plus Antibacterial suture was statistically significantly larger than the BSS groups.

## Discussion

Since Ethicon gained approval to market their antimicrobial polyglactin 910 with triclosan sutures in 2002, their reception has been mixed with recent trials and meta-analyses providing evidence of their benefit (1–9), despite individual studies initially failing to show a reduction in SSIs (10, 11). In vitro studies of AMS have provided further support for their benefit. Rothenburger et al. used zone of inhibition assays to demonstrate that polyglactin 910 with triclosan sutures inhibit the growth of S. aureus and S. epidermidis even after passes through tissue and aqueous immersion (14). Edmiston et al. demonstrated decreased adherence of Gram positive and negative bacterial species to AMS which lead to decreased local bacterial loads for at least up to 96 hours (15). This latter finding could help explain the findings of a study by Ford et al. that reported less postoperative pain in general pediatric surgical patients in which AMS were used, although interestingly it failed to show a concomitant decrease in SSIs within the group (16).

Most initial studies have investigated the utility of AMS in GI procedures, but recent studies have begun to explore broader applications. Rozzelle et al. showed decreased CSF shunt infection rates with the use of AMS (8). Others have demonstrated decreased bacterial loads around implant sites in rat models, suggesting a role for AMS in decreasing implant infections (13). With these added applications, others have attempted to develop sutures for ophthalmological procedures (12).

Despite the increased upfront costs, AMS have demonstrated their cost effectiveness by decreasing overall costs related to the management of wound infections. One study estimated the widespread use of AMS at one center could save approximately $1.5 million in a year by avoiding costs related to SSIs (3). Additionally, another study found AMS to significantly reduce the incidence of SSIs and subsequently lead to reduced overall costs as its analysis found the average cost of SSI management to be $2,310 (7).

Given the reduction in SSIs and subsequent associated costs, our study sought to replicate the antimicrobial activity of marketed AMS at a reduced overhead by having the surgical technician place the sutures in a bacitracin solution that is prepared at the beginning of the procedure for use as irrigation in lieu of directly placing them on the table. An example of this procedural change is shown in Figure 1. Our in vitro antimicrobial zone of inhibition assays suggests that our intraoperatively prepared Ethicon 0 VICRYL® BSS provide superior antimicrobial activity compared to unsoaked Ethicon 0 VICRYL^®^ suture. This benefit is potentially due to a Bacitracin coating that forms during the soaking process, evinced by sustained antimicrobial activity even after allowing 6 hours of drying. Yet, our study also suggests that while the soaking process confers some antimicrobial activity, it fails to retain the full antimicrobial exhibited by an aliquot of the Bacitracin irrigation solution itself. Additionally, the current AMS on the market—Ethicon’s Coated 0 VICRYL^®^ Plus Antibacterial suture— exhibits superior antimicrobial activity to our intraoperatively BSS, likely due to enhanced drug-eluting properties related to their manufacturing process. However, as zones of inhibition signify drug potency, the differing sizes may not be clinically relevant. The BSS displays a consistent and robust antibacterial activity, with the Ethicon Coated 0 VICRYL^®^ Plus Antibacterial suture demonstrating a greater drug concentration and eluting property as evinced by the greater zone of inhibition sizes. Therefore, given local potency of the BSS in preventing bacterial growth, there is doubtful to be clinically meaningful differences in SSI incidence using either suture. Further studies examining the antimicrobial activity of our BSS after passes through tissue and aqueous immersion could help identify the longevity and overall viability of this antimicrobial coating in a clinical setting.

Ultimately, this study suggests that with the minor procedural change of placing sutures in a bacitracin irrigation solution intraoperatively instead of directly on the table, it can achieve some of the antimicrobial effect of the AMS currently on the market. This may result in reduced rates of SSIs and associated costs without major procedural change and at reduced overhead. Future studies should be undertaken utilizing this change to fully ascertain its clinical efficacy and price-reducing effectiveness.

## Conclusions

AMS lower the incidence of SSI and subsequently decrease associated costs despite increased overhead cost. We attempted to develop a novel method that could reproduce the antimicrobial benefits while lowering these overhead costs. Our zone of inhibition assays suggests that our method of intraoperatively soaking sutures in the antibiotic irrigation confers antimicrobial activity against S. aureus and MRSA that is superior to plain suture, yet inferior to the AMS currently on the market. Further studies are needed to determine if our BSS can better replicate the antimicrobial activity of the AMS currently on the market and if the antimicrobial activity observed in our study has clinical implications.

